# What stabilizes pre-folded structures in the intrinsically disordered α-helical binding motifs?

**DOI:** 10.1101/2022.01.28.478151

**Authors:** San Hadži, Samo Purič, Uroš Zavrtanik, Wim Vranken

**Affiliations:** Department of Physical Chemistry, Faculty of Chemistry and Chemical Technology, University of Ljubljana, 1000 Ljubljana, Slovenia; Graduate Study Program, Faculty of Chemistry and Chemical Technology, University of Ljubljana, SI-1000 Ljubljana, Slovenia; Artificial Intelligence Laboratory, Vrije Universiteit Brussel, Pleinlaan 2, 1050 Brussels, Belgium; Interuniversity Institute of Bioinformatics in Brussels, ULB/VUB, Triomflaan, 1050 Brussels, Belgium; Structural Biology Brussels, Vrije Universiteit Brussel, Brussels 1050, Belgium; VIB Structural Biology Research Centre, Brussels 1050, Belgium

**Keywords:** intrinsic disorder, pre-folding, folding-upon-binding, recognition motif, dataset

## Abstract

Many examples are known of regions of intrinsically disordered proteins (IDPs) that fold into α-helices upon binding their globular protein targets. In their unbound state these regions possess a small amount of residual helicity, referred to as pre-folded structure, which has been studied on case by case basis. In order to investigate what determines these pre-folded structures we compiled a database of peptides that fold-upon-binding, and experimentally characterized their helicity in the unbound and target-bound state. These regions are more hydrophobic and lack proline residues compared to IDPs in general. On average they possess about 17% helicity in the pre-folded state and gain 40% of helicity upon target binding. We observe that the locations of pre-folded helical regions strongly overlap with those in the target-bound IDPs. To understand this correlation, we analyzed per-residue energetic contributions stabilizing helical structure and found that target-interacting IDP have higher helix propensity. Notably, leucine is the most common residue involved in IDP-target interactions and, due to its high helix propensity, it strongly stabilizes pre-folded helical structures. For many IDP binding motifs, particularly those enriched in leucine, we observe that they not only mediate target-interactions but also confer stability to the pre-folded structure. Collectively, this shows that the formation of pre-folded helical elements is coupled to the IDP-target interactions, explaining why such elements are a common feature of α-helical binding motifs. Moreover, it probably explains the preference for leucine among IDP-target hotspots, even though this residue is underrepresented among hotspots in the interfaces between globular proteins.

## Introduction

Intrinsically disordered proteins are characterized by a lack of stable three-dimensional structure and a specific amino acid composition, enriched in charged and depleted in hydrophobic residues [1]. Nevertheless, they perform key physiological functions and are often associated with many essential cell processes, such as transcription regulation, signal transduction, assembly of multiprotein complexes, and formation of membrane-less organelles [2,3]. It is estimated that about 25-30% of eukaryotic proteins are completely disordered and an even larger fraction contain long disordered regions [4]. Central to their function are interactions with globular proteins, where IDPs often adopt a well-defined confirmation in their bound state or can equally remain dynamic and disordered [5].

Among different types of interactions between IDPs and globular proteins, the most prevalent and best studied class represents IDPs that fold into *α*-helices upon binding the target [4]. These are particularly common among the eukaryotic transactivation domains (reviewed in [6]). Interaction affinities of IDPs that fold upon binding are typically in the high nanomolar to micromolar range, however examples of ultra-high affinities in the picomolar rage have also been measured [7,8]. While it was initially suggested that these IDPs are largely disordered in the unbound state [9,10], later observations indicated such regions may exhibit substantial amounts of residual helical structure [11]. These regions have been referred as molecular recognition elements, pre-structured motifs, pre-folded or preformed elements, transient or residual secondary structures [12]. An early bioinformatics study has found that the residual structure may coincide with the structural elements in the final, target-bound IDP structure, suggesting that the unbound IDP structure predisposes the bound-state conformation [13]. The mentioned study was however based on a relatively small number of systems, as few studies were available at the time. A more recent survey of experimentally studied IDPs in their unbound state listed over 50 examples where IDPs were shown to possess a significant amount of secondary structures in the unbound state, but did not systematical analyze to what extent the residual structure coincides with the target bound one [11]. It has been commonly argued, based on simple thermodynamic arguments, that such pre-formed elements should lower the entropic penalty of IDP folding and thereby increase the IDP-target affinity [14]. However, while this seems to be true for some cases [15,16], not all studies point in this direction [17]. Similarly, different results have been observed in kinetic studies, where residual helicity in some cases affected the rate constants, while in other systems it has no substantial effect [18,19]. Collectively, the pre-structured helical elements have been observed in many systems, but their functional importance appears to depend on the IDP-target system.

In order to understand how much residual structure is generally present in IDPs, we compiled a database of residual helicities of peptides corresponding to α-helical regions of IDPs, experimentally determined by circular dichroism. The average helix content in the unbound state is around 15% and increases on average by 40% upon target binding. Using a combination of NMR data and helical predictions from AGADIR, we show that the positions of helical regions in the unbound IDP state are in general strongly correlated with the locations of helices that form upon target binding. Analysis of energetic contributions stabilizing helical conformation shows that the residues involved in IDP-target interactions, in particular leucine, also confer strong stabilization of helical structure in the unbound state. The high occurrence of leucine among IDP-target hotspot interactions, which is otherwise depleted in the interactions between globular proteins, is most likely related to the mentioned stabilization of pre-folded structures and thus typical for α-helical binding motifs. Thus, IDP-target interactions appear to be coupled to the formation of pre-formed helical elements, which explains the ubiquity of pre-folded structure among the intrinsically disordered α-helical binding motifs.

## Results

### A database of experimental helicities of the α-helical binding motifs in the unbound state

The formation of pre-folded structure (also known as residual or transient helical structure) in intrinsically disordered α-helical binding motifs has been studied on a case by case basis for many systems. In order to systematically examine the extent of pre-folded structure in these systems, we compiled a database of experimentally determined helicities derived from circular dichroism spectra using published data complemented with our own measurements. The database consists exclusively of IDPs, or segments thereof, that form α-helices upon binding their partner proteins, and does not include coiled-coils or small globular proteins that fold conditionally. The experimental helix content of these unbound IDPs was determined from the CD spectra as described in *Methods.* Briefly, literature CD spectra were digitalized to obtain the intensity at 222 nm, which is proportional to the number of peptide bonds in the helical conformation. Using molar ellipticity of pure helix and coil conformation [20], we estimate the fraction of residues in helical conformation i.e. fractional helicity. In this manner we analyzed 53 IDPs form the literature and 12 IDPs whose CD spectra were measured in our laboratory, resulting in a dataset of 65 IDPs (Table S1, Fig. S1). The average length of IDP peptides is 30 residues. In majority of cases the CD spectra were acquired at the room temperature (20 or 25 °C) and at physiological conditions (pH 7-7.5, ionic strength 0.1 M), while in 10 cases the temperature was 4 or 5 °C. Additionally, we cross-checked the BMRB database for the NMR chemical shift data, which were available for 25 of IDPs included in the database (Table S1). While the CD data provide the average helix content, the NMR chemical shift measurements can be used to analyze the IDP helicity on the per residue basis. Finally, a search in the protein data bank (PDB) provided the structures of these IDPs in their target-bound state for 57 of these IDPs, while for the remaining 8 IDPs evidence that they fold into α-helical structures comes from other data *(e.g.* CD). Table S1 lists the names of IDPs and their UNIPROT codes, sequence of the IDP segment used for CD measurement, experimental conditions, experimental helicity, BMRB entry for the free IDP, PDB code for the target-bound IDP, and additional information.

### Sequence characteristics of intrinsically disordered α-helical binding motifs

To characterize the sequence properties of the IDPs from the dataset we first analyzed their net charge and hydropathy. The charge-hydropathy plot shows that intrinsically disordered α- helical binding motifs cluster near the border commonly assumed to separate globular and disordered proteins (Fig. 1A) [1]. The average mean hydrophobicity of the dataset is 0.42 on Kyte-Doolittle scale and the average absolute charge is 0.12, 62% IDPs are negatively charged. Overall, it appears that the sequence characteristics of intrinsically disordered α- helical binding motifs differs from that of IDPs in general, which are usually more charged and less hydrophobic [1]. To understand these differences in more detail we analyzed the amino acid composition of our dataset and compared it to the one from globular proteins [21]. Our IDP sequences appear to be enriched in all charged residues (D, R, E, K), two polar (Q, N) and hydrophobic (M, L) ones, but strongly depleted in other hydrophobic and aromatic residues (I, V, F, Y) as well as T, P, C (Fig. 1B). At first glance, enrichment of charged and depletion of aromatic and hydrophobic residues resonates with the general sequence composition of IDPs. However, there are important differences in the composition profiles between α-helical binding motifs and IDPs in general. For example, α-helical motifs are depleted in P and S, whereas these two are among the most enriched amino acids in IDPs [1,22]. Furthermore, the L is strongly enriched in α-helical motifs, but depleted in IDPs. The composition profile for E, M, K, A is similar for both groups of IDPs, while α-helical motifs have higher contents of I and W (they are more depleted in IDPs) and of charged residues D, R [1]. These compositional differences can be rationalized by considering two aspects: helix folding and the formation of IDP-target interactions. Depletion of proline, a strong helix breaker, likely facilitates folding into α-helices upon target binding. On the other hand, a preference for the hydrophobic and aromatic residues I, L and W is probably related to the formation of hydrophobic IDP-target interfaces. Enrichment of charged residues D and R, relative to general IDP sequences, perhaps helps to compensate for the rather high hydrophobicity of these peptides, so maintaining disorder in their unbound state. Using a different reference set for globular proteins, results in somewhat different amino acid frequencies, but the qualitative differences discussed above remain similar (Fig. S2).

**Figure 1.**
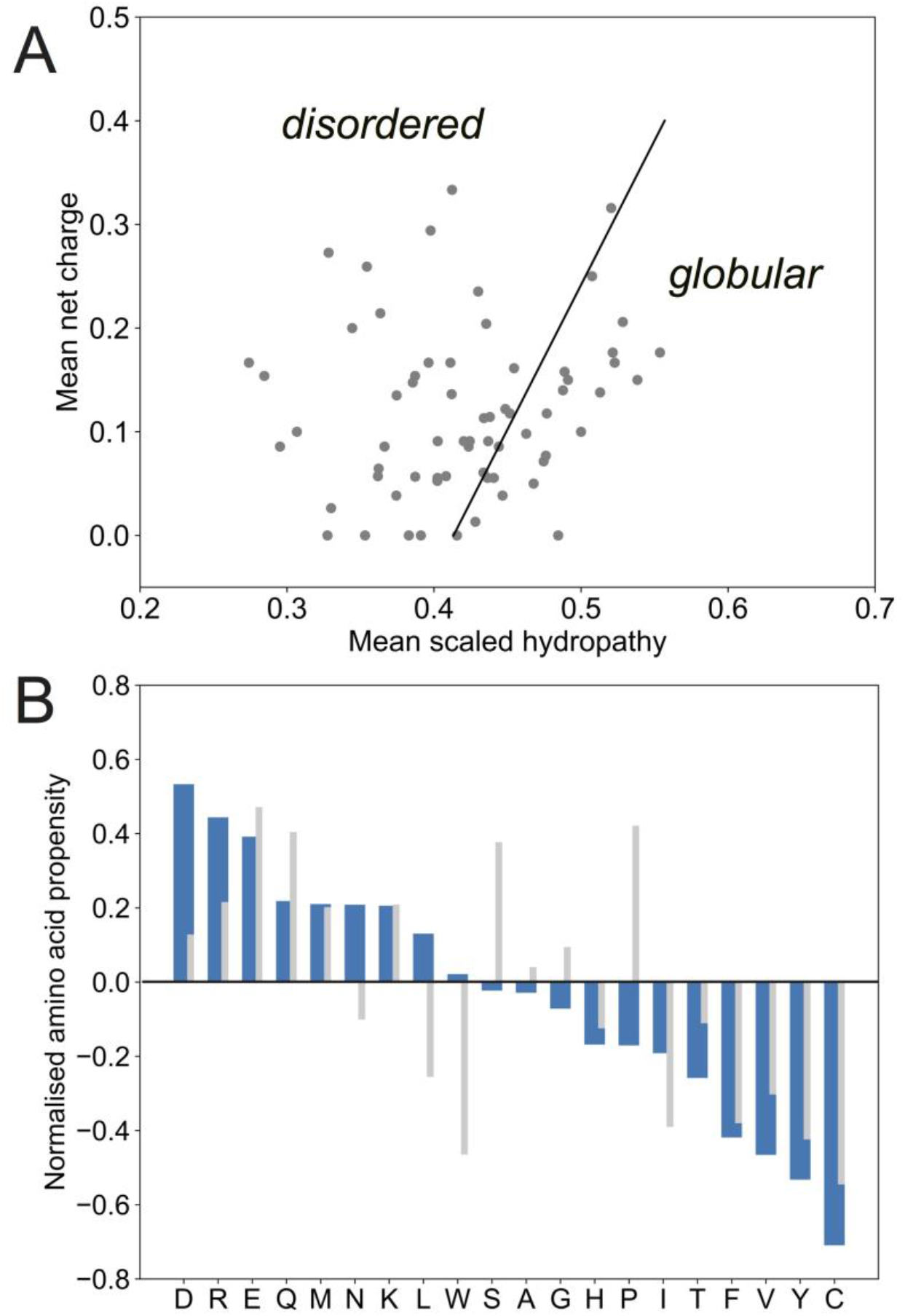
Sequence characteristics of intrinsically disordered *α-*helical binding motifs. **A.** The charge-hydropathy plot shows that majority of intrinsically disordered α-helical binding motifs in our dataset cluster around the border (black line) that separates globular proteins from the intrinsically disordered proteins [1]. **B.** Amino acid composition of disordered α-helical binding motifs from the dataset is normalized to the frequencies observed in globular proteins [21] is shown by blue bars. Positive values indicate that these amino acids are enriched relative to the globular proteins, while negative values indicate amino acid depletion. Gray bars show the corresponding amino acid composition of IDPs from the Disprot database (as reported in [22]).

### Distribution of IDP helicities in the unbound and target-bound state

The distribution of overall peptide helicities for unbound IDPs as estimated from the CD signal intensity is shown on Fig. 2A (blue bars). The average residual helicity of α-helical IDPs in the dataset is 17%, with the majority of values between 0 and 30%. Four IDPs show helicities that are higher than 30%: BMF with 67 % helicity (UNIPROT: Q91ZE9, residues 124-158), NOXA_B with 51 % (UNIPROT: Q9JM54, residues 64-98), FCP1 with 35 % (UNIPROT: Q9Y5B0, residues YG944-958) and PaaA2 with 37 % helicity (UNIPROT: Q8XAD5, residues 35-63). On the other hand, for three quarters of the IDPs in our dataset the helix content is lower than 10 %. In order to estimate the extent of folding upon binding the target, we analyzed the target-bound IDP conformation from the PDB structure (Table S1). The number of residues that adopt helical conformations was determined based on their torsion angles, and then used to calculate fractional helicity in the target-bound state (see *Methods).* While in most cases the peptide segment used for CD measurement corresponded to the one observed in the target-bound structure, these were sometimes longer compared to those in the PDB file. In this case it was assumed that the residues which are not seen in the target-bound experimental structure remain disordered. Thus, the target-bound helicity is defined with respect to the IDP sequence in the unbound state characterized by CD. In contrast to the rather uniform distribution of helicities in the unbound state, the target-bound IDPs show a very broad distribution of helicities ranging from 14 % to 100 %, with the average helicity around 60% (Fig. 2A, orange bars). Thus, in our dataset intrinsically disordered α-helical binding motifs fold to different degrees with the average increase in helicity around 42%. Interestingly, values of unbound and bound helicities are not correlated (Fig. S3).

**Figure 2.**
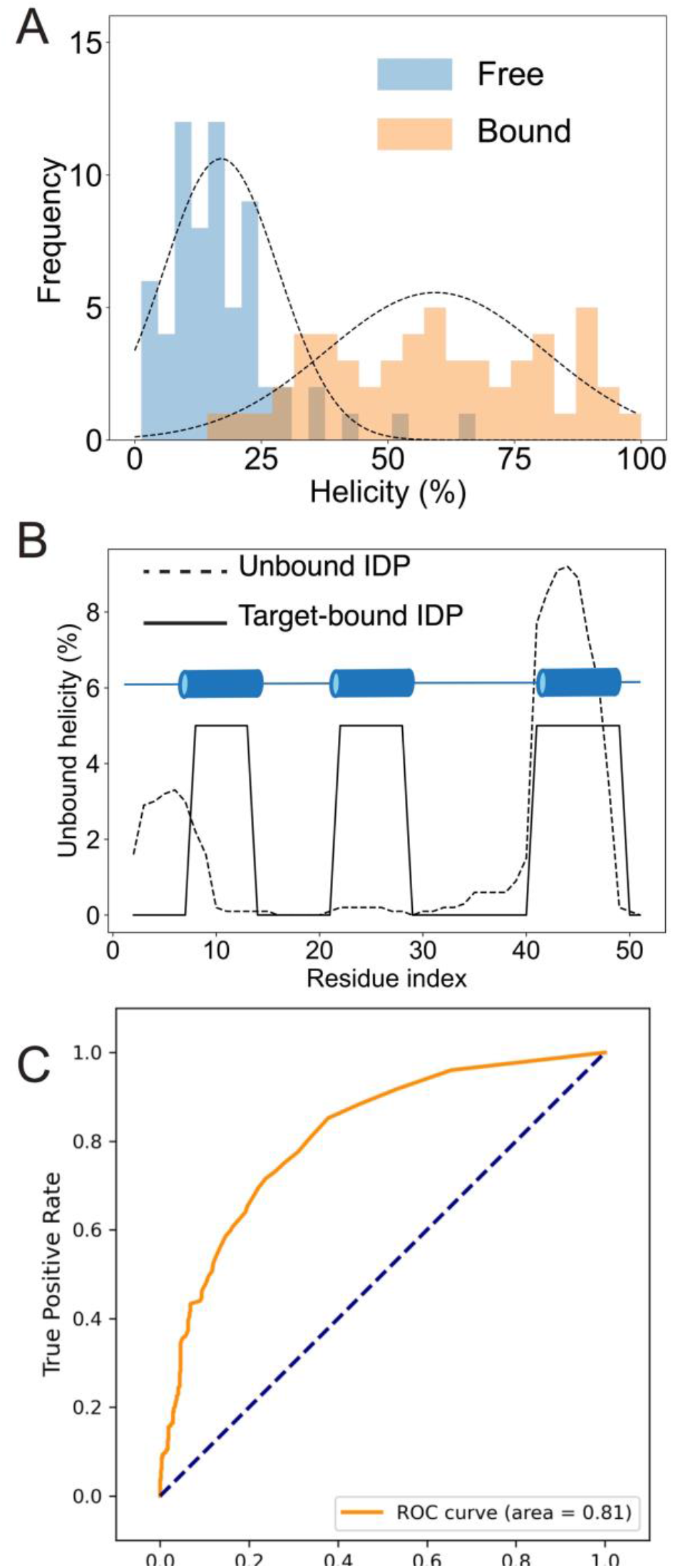
Helicities of intrinsically disordered *α-*helical binding motifs in their unbound and target-bound states. **A.** Distribution of helicities in the unbound (blue) and target-bound state (orange). The unbound helicities are calculated from the CD data, while target-bound helicities are calculated from the structures of IDP-target complex as described in Methods. **B.** An illustration of overlapping helical regions for the Hif1-α transactivation domain. The dotted line shows the helicity in the unbound (pre-folded) state as calculated with AGADIR, while the solid line shows helical elements in the target-bound structure (having 0 or 100% helicity). **C.** ROC curve of AGADIR predicted per-residue helicity for the IDP peptides in our dataset compared to the helical positions observed in the target-bound structures shows that helix positions in the unbound and target-bound state are strongly overlap.

### Helical elements in unbound IDPs overlap with the helices in the target-bound state

While the analysis of CD spectra provides an experimental estimate of the average helix content within the peptides, it provides no information on the locations of the helical regions in the IDP sequence. To investigate the local formation of pre-folded helical structure, we used different sequence-based and NMR chemical-shift based helix prediction tools. For sequence-based tools we used AGADIR, which implements the helix-coil theory for prediction of helical content in peptide [23] and has been parametrized on a large sample of CD and NMR data, and the b2btools suite [24], which includes DynaMine for backbone dynamics[25] and EFoldMine for early folding propensity [26]. For the chemical-shift based tools, which estimate the per-residue conformational population directly from the NMR chemical shift values observed for the peptides in solution, we used δ2D [27] and cheSPI [28]. An example for the predictions generated using these tools is shown on Fig. S4 for one of the PaaA2 peptides.

We next investigated to what extent the helical elements in the unbound state overlap with those observed in the IDP-target complex. An example of the distribution of helices in the unbound state as predicted by AGADIR and those in the target-bound structure is shown in Fig. 2B for the Hif1-α IDP, where the partial overlap between the two helical regions is clearly observed. A more systematic analysis of such overlaps for all IDPs in the database in terms of ROC curves is shown in Fig. 2C. In this graph, a sliding cutoff is used on the per-residue helicities predicted by AGADIR to classify residues into helical or not helical, and these classes are then compared to the ground truth from the PDB structures (represented by the orange line in Fig. 2C). If there is a perfect match between prediction and experiment, the orange line goes straight to the top left corner, with a true positive rate of 1.0 and a false positive rate of 0.0, whereas a fully random prediction follows the dotted diagonal line. We find that pre-folded helices, as obtained from the AGADIR prediction, have a ROC area under the curve (AUC) of 0.82, which shows the AGADIR predicted helical propensities correspond well with the locations of target-bound helices. A comparable performance of 0.80 of ROC AUC is obtained by combining the b2bTools predictions of early folding and helix propensity, indicating peptide regions with helical propensity and a tendency to fold independently (Table S2). The chemical shift based values are dependent on the method: the δ2D-estimated helix population is somewhat predictive with a value of 0.63, but the beta population is strongly negatively correlated (0.21 ROC AUC, which corresponds to 0.79 when the values are reversed), indicating a profound lack of extended conformation in the peptides (Table S2). The cheSPI method, on the other hand, identifies helix and sheet propensity to both be predictive at values of 0.70 (Table S2), showing that the information is present in the experimental data, but that the cheSPI interpretation in terms of conformation is quite different from δ2D. Collectively, these results show a strong correlation between the predicted helix position in the pre-folded elements with the target-bound IDP helix conformation, while the interpreted chemical shift values correlate with the final bound conformation, but that the interpretation thereof is method-dependent.

We also observed that the overall helix content predicted by AGADIR seems to give systematically lower average helicity values compared to those obtained for the isolated peptides from CD. Whereas the experimental average helicities are around 17 %, the average of AGADIR predictions is significantly lower, about 4 % (t-test, *N_1_=N_2_=65, p*<10^-12^, Fig. S5). Nevertheless, AGADIR and CD helicities correlate (R=0.79, slope=0.16) at least for the data acquired in our laboratory (Fig. S5). It is difficult to speculate why AGADIR systematically predicts lower absolute helicities of IDPs. AGADIR has been parametrized on a CD and NMR helicity data consisting mainly of the alanine-rich peptides, which significantly differ from the peptides studied here. The presence of aromatic residues can contribute to CD intensity and effect the estimation of helicity [29]. The aromatic contributions to the CD signal are sitespecific and depend on the orientation of sidechains therefore it is difficult to apply these corrections in simple terms. In our dataset there are 25 cases where an aromatic residue (Y or W) is present at the non-terminal positions (terminal positions lead to negligible effects). It has been shown that the aromatic contributions tend to underestimate the helicity by <5%, which would even enhance the discrepancy with AGADIR predictions [30]. The observation that AGADIR underestimates absolute helicity for IDPs has also been noticed previously [19]. Collectively, these results show that while AGADIR appears to correctly predict helical regions in a relative sense – it can identify residues more likely to be helical – it does underestimate the absolute values of helix content in IDPs. This is important, since AGADIR has often been used in the literature to asses helicity of the IDP regions.

### Analysis of IDP residues mediating interactions with the target

We next investigate why the locations of helical elements in the unbound state coincide with those in the target-bound conformation. A plausible hypothesis could be that the IDP residues mediating target-interactions also stabilize helical conformations in the pre-folded state. In other words, the formation of pre-folded elements could be coupled to target interactions. To investigate this hypothesis further, we analyzed IDP residues mediating target interactions and classified them as target-contacting or non-contacting based on the sidechain solvent accessibility in the IDP-target complex (see Methods). Additionally, we also introduced a group of IDP hotspot residues, which are the residues that form the energetically most favorable IDP-target interactions (see Methods). We find that hydrophobic and aromatic residues strongly dominate the contacting IDP positions, whereas polar and charged residues are favored at non-contacting positions (Fig. 3). Over 55% of contacting residues are hydrophobic and majority of those are represented by L (>20%). A strong preference for contacting over non-contacting positions is also observed for the residues I, V, M and F. In contrast, charged amino acids are preferred on the non-contacting positions at approximately similar frequencies, around 10% (Fig. 3A). Among polar residues S is the most common residue on non-contacting positions (10% frequency) and has high preference over contacting positions.

**Figure 3.**
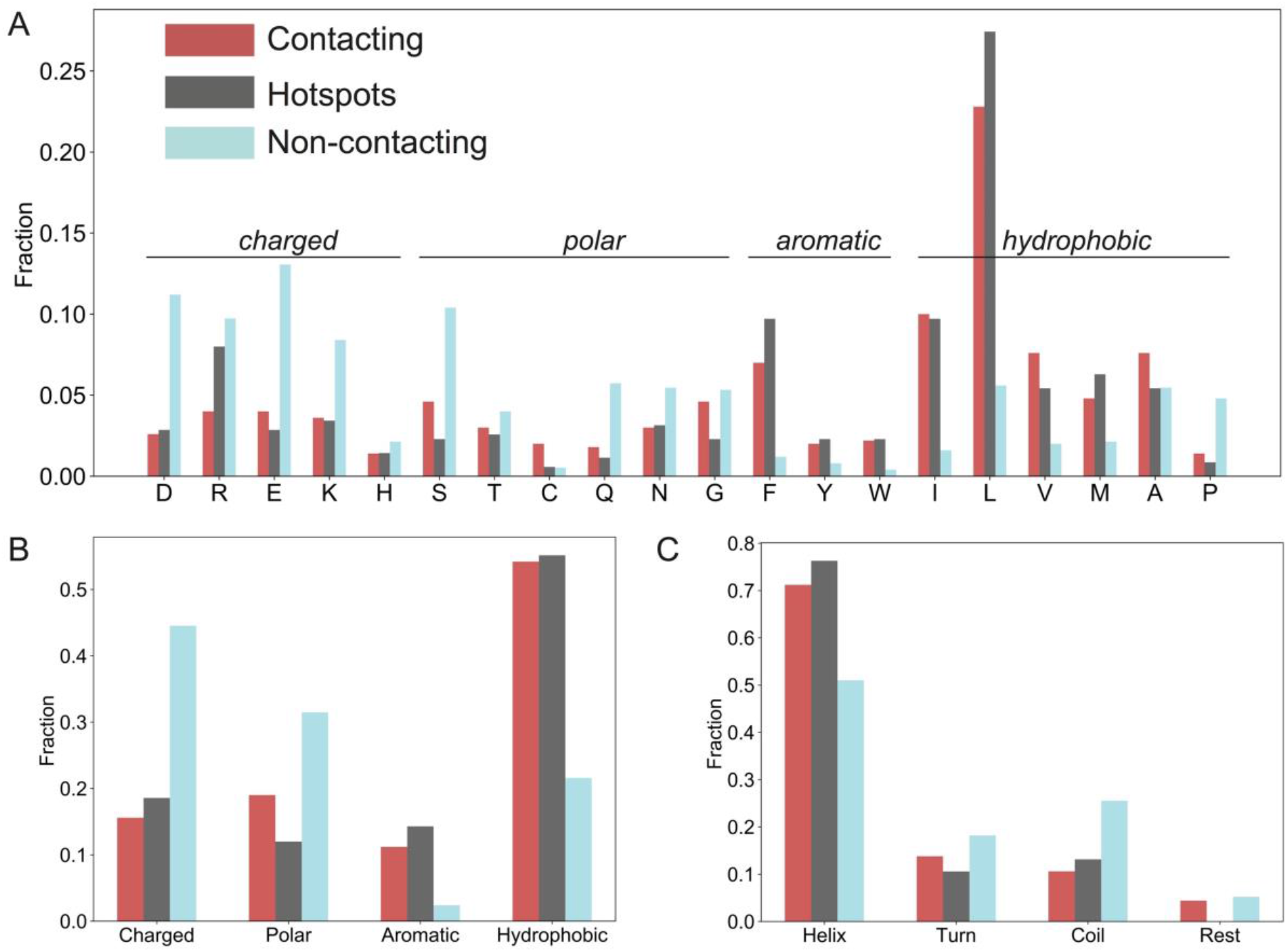
Analysis of target-bound *α-*helical binding motifs. **A.** Amino acid frequencies for the contacting, non-contacting and hotspot IDP residues from the IDP-target complexes. A strong preference for leucine as a contacting and hotspot residue is immediately apparent. Bars show the frequencies of contacting, non-contacting and hotspot residues for different amino acids groups (**B**) and their occurrence in different secondary structures (**C**).

We observe that vast majority of hotspots residues adopt helical conformation in the target-bound state (>75%) while around 10% are found in coil conformation (Fig. 3C). Around 10% of hotspots are in turn conformation, which in most cases corresponds to the α-turn observed at helix termini due to helix fraying. This supports the idea that residues forming hotspot interactions adopt helical conformation in the target-bound conformation and thus drive helix folding. As shown above the hotspot residues are already pre-arranged in a significant proportion prior to binding, given the findings of the predictions (Fig. 2). The residue that makes the most favorable interactions (hotspots) is L, which is involved in 27% of all hotspots, while all hydrophobic residues make over 55% of all hotspots. The second most common hotspot residues are isoleucine and phenylalanine (each representing 9.7% of hotspots). To show which residues preferentially form hotspots we normalize the hotspot frequency by the contact frequency (Fig. S6). For example, arginine is not frequently contacting residues, but when it does form contact it preferentially forms a hotspot interaction (Fig. 3A, S6). Such a preference to make hotspot interaction is typical for R but also observed for F, Y, L and M (Fig. S6). The preference of Y and R to preferentially establish hotspot interactions has also been observed in antibody-antigen interactions, whereas in protein-protein interactions R, Y, and W are the most common hotspots [31,32]. Strikingly, leucine is depleted as a hotspot residues in the interfaces of globular proteins [31,32]. It therefore appears that the high occurrence of leucine hotspots is a distinctive as well as defining feature of interfaces formed by the intrinsically disordered α-helical binding motifs.

### Target-interacting IDP residues have high helix propensity and stabilize pre-folded structure

In order to elucidate the conformational determinants of the pre-folded elements, we analyzed the energetic contributions that stabilize helical conformation. Previous studies have established that energetics of helix folding can be separated into contributions related to helix nucleation, helix propagation, sidechain and capping interactions [33,34]. Amino acids have a different intrinsic helix propensity Δ*G*_CH_, which is the per-residue free energy for a transition from coil to helix conformation and depends on the nature of the sidechain. The Δ*G*_CH_ values have been determined for all amino acids and are commonly expressed in terms of helix propensity scales [35]. For example, alanine favors helical conformation and has the highest intrinsic helix propensity, therefore the negative Δ*G*_CH_=-250 cal/mol. In contrast, glycine is a helix breaker (Δ*G*_CH_=+750 cal/mol) [35,36]. For each IDP peptide we calculated the average per-residues helix propensity for all residues and also for contacting, non-contacting and hotspot residues (Fig. 4A). Overall, the average helix propensity for all IDP residues in the dataset is positive (Δ*G*_CH_=260 cal/mol per residue), meaning that IDPs are mainly unfolded in their unbound state (Fig. 4A). To put this number in perspective we calculated the average Δ*G*_CH_ for randomly selected sets of peptide sequences of different lengths found in the Disprot database [37] resulting in the average value Δ*G*_CH_=360 cal/mol (Fig. S7). Thus, intrinsically disordered α-helical binding motifs have more favorable per-residue helix propensity compared to general IDPs in the Disprot database (t-test, *N_1_=57, N_2_=2000, p*<10^-17^). More significantly, we found that the stabilization of helical structure is not equally distributed among all residues. Target-contacting residues in our dataset have more higher helix propensity (Δ*G*_CH_=170 cal/mol, Fig. 4A) compared the average of all residues in the sequence (t-test, *N_1_=N_2_*=57, *p*=10^-4^). Contacting residues that establish hotspot interactions (hotspot residues) have even higher helix propensity (Δ*G*_CH_=140 cal/mol, t-test, *N_1_=N_2_*=57, *p*<10^-7^, Fig. 4A). In contrast, helix propensity of non-contacting residues does not differ significantly from that of the whole IDP sequence (Fig. 4A). These results show that contacting residues and hotspots in particular have a higher propensity to form helices. This result is most likely related to the strong representation of leucine in the group of contacting and hotspot residues, which has the highest helix propensity among amino acids besides alanine.

**Figure 4.**
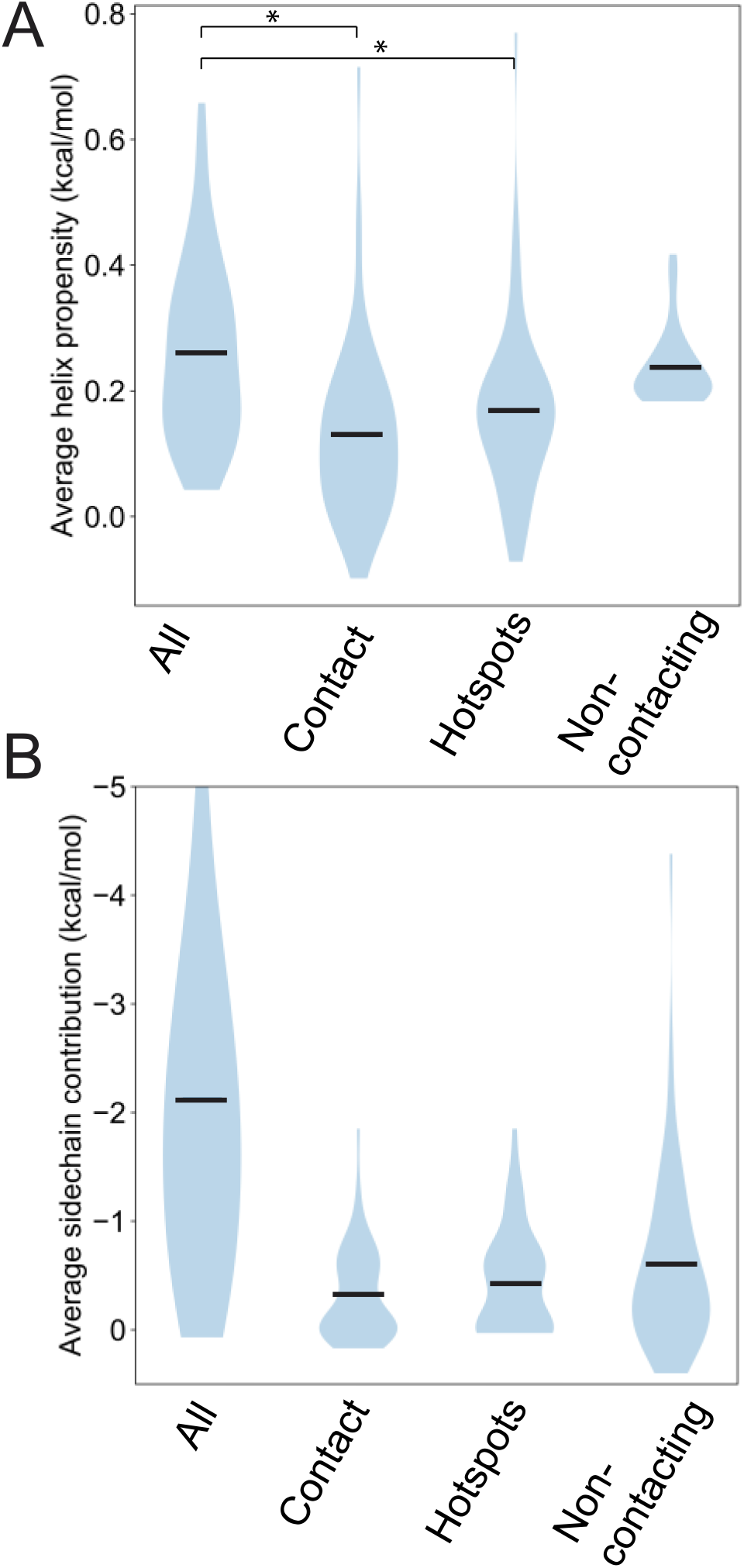
Target-interacting IDP residues also stabilize pre-folded helical structure. **A.** The intrinsic helix propensity (Δ*G*_CH_) corresponds to the per-residue free energy accompanying coil-helix transition. Using known values of Δ*G*_CH_ for different amino acids [35] we calculate the average per-residue Δ*G*_CH_ for the whole IDP sequence, as well as for residues that contact the target, those that make hotspot interactions and the non-contacting residues. The average Δ*G*_CH_ for the contacting and hotspot residues is significantly lower compared to the whole sequence, meaning that these residues stabilize helical structure. **B** The average contributions of intrahelical sidechain interactions to helix stabilization conferred by all residues (whole IDP sequence), contacting, hotspot and non-contacting residues, respectively. The sidechain interactions mediated by contacting and non-contacting residues both stabilize prefolded helical structure. Only a small number of hotspot residues makes a significant (−0.32 kcal/mol on average) contribution via sidechain stabilization.

Having established that IDP hotspot residues have higher helix propensity compared to the rest of the IDP sequence, we next investigated the energetic contribution of the intrahelical sidechain interactions in stabilization of helical elements. Experimental studies on model peptides have identified that sidechains from the *i* and *i*+4 (or *i*+3) residues can significantly contribute to helix stability [34]. We observed that in our IDP dataset there are on average 5 such intrahelical sidechain interactions per IDP that affect helix stability (Fig. S8). On average all these interactions contribute about −2.1 kcal/mol to helix stabilization per IDP (Fig. 4B). The majority of these interactions (>55%) are between oppositely changed residues *(e.g.* E-K bridges between *i-i*+4 residues), which are formed by the residues at the non-contacting positions that are also part of the helix. The second most common type are interactions between hydrophobic sidechains (25%) mediated by the contacting and hotspot IDP residues (Fig. S9). Intrahelical interactions mediated by the non-contacting residues (which are part of helix in the target-bound state), contribute around −0.64 kcal/mol while those from contacting ones about −0.43 kcal/mol (Fig. 4B). Interactions mediated by hotspot residues stabilize helices by −0.32 kcal/mol on average (Fig. 4B). Collectively these data show that contacting and hotspot residues stabilize helical structure due to their increased helix propensity as well as favorable intrahelical sidechain interactions. Non-contacting residues that are part of the helix also stabilize helical structure by sidechain interactions, but they do not have increased helix propensity. Overall, stabilization of helices by target-interacting IDP residues is most likely due to high frequency of leucine, which has both high propensity and can establish favorable *i-i*+4 sidechain interactions. Thus, hotspot residues are not only important for target interactions but are also likely to at least transiently stabilize pre-folded elements through favorable side-chain interactions.

### Leucine-rich binding motifs confer stability to pre-folded elements

Prompted by the observation that target-binding IDP residues also stabilize the prefolded structures, we investigated to what extent common α-helical binding motifs might stabilize pre-folded IDP structures. For example, a very common IDP binding motif encountered in the intrinsically disordered transactivation domains is the hydrophobic LXXLL motif, with X being any amino acidic. The effect of leucine residues in this motif in promoting helicity is shown on Fig. 5A. The host sequence, for which we use a repeated SETSTTSSS sequence having the same average helix propensity Δ*G*_CH_=+260 cal/mol as observed for IDPs in our database, is disordered and shows no tendency for the pre-folded α-helical structure (solid line, Fig. 5A). The presence of three consecutive leucine residues strongly increases IDP helicity as calculated by AGADIR due to the high intrinsic helix propensity of leucine residue (dashed line, Fig. 5A), but not due to sidechain interactions. When leucine residues are placed at appropriate spacing (e.g. *i-i+3 or i-i+4*), so enabling formation of favorable sidechain interactions, the helicity increases even further (dotted line, Fig. 5A). These calculations illustrate that the presence of a leucine-rich binding motif not only leads to favorable IDP-target contacts, but also promotes helical structure. In contrast, presence of other hydrophobic residues such as isoleucine or valine, which could equally mediate IDP-target contacts but have lower helix propensity, does not significantly stabilize pre-folded helical structure (Fig. S10).

**Figure 5.**
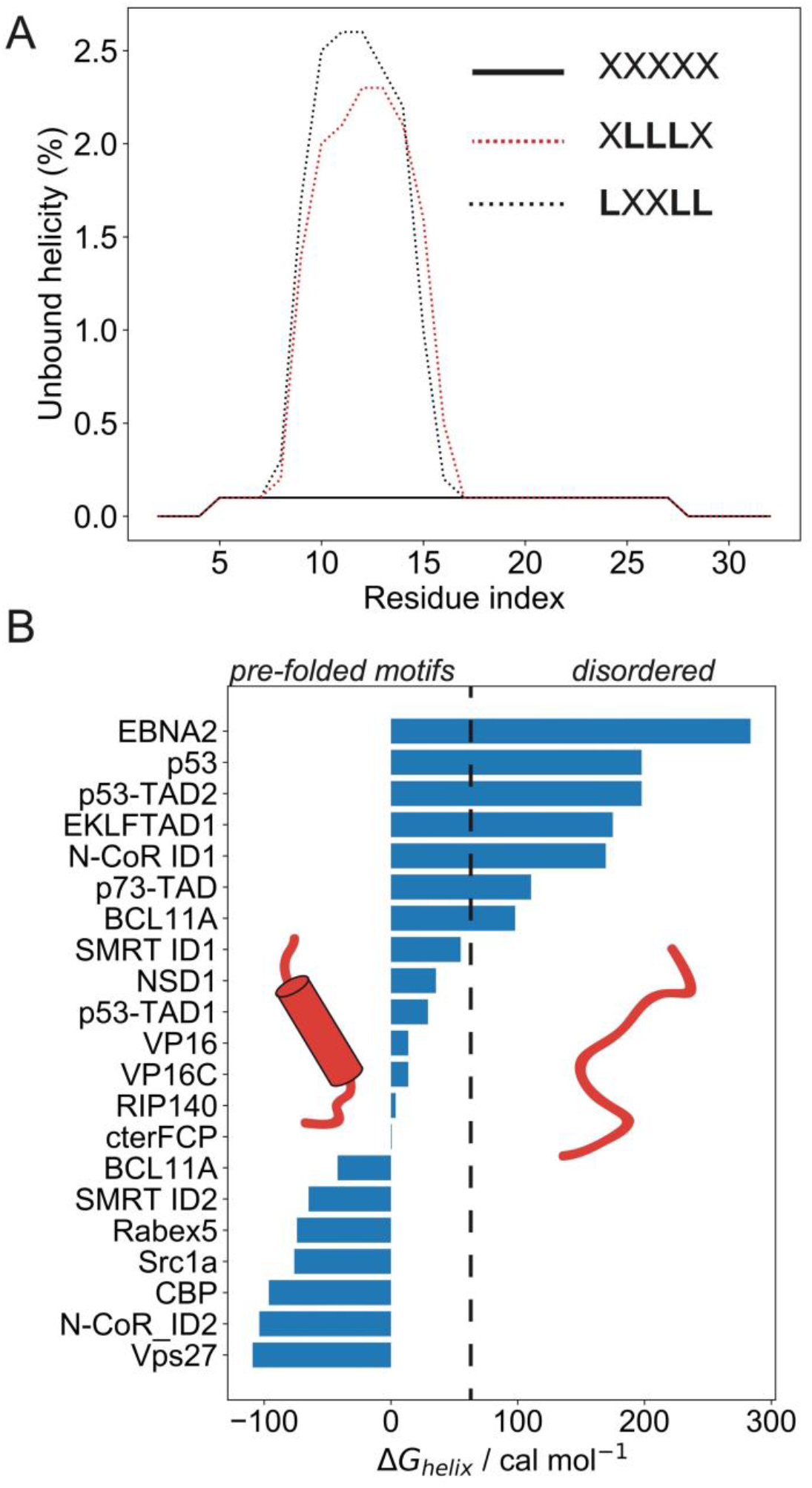
Leucine residues in the IDP binding motifs confer stability to the pre-folded helical structure. **A.** AGADIR calculated helicity shows the effects of leucine residues, which are frequent among target-interacting residues, on the unbound IDP helicity. A random host sequence with IDP characteristics has low helicity as shown by the black solid line. Addition of few leucine residues, considerably increases helicity due to high helix propensity of leucine and stabilizes pre-folded helical elements (dotted black line). A common interaction motif LXXLL has even higher helicity due to intrahelical sidechain interactions that further stabilize pre-folded helical structure (red dotted line). **B.** Bars show the average value of helix stabilizing in some common IDP binding motifs. The average per-residue helix stabilization is calculated as the sum of intrinsic helix propensity (Δ*G*_CH_) and sidechain contribution for each motif. The dotted line separates motifs that promote pre-folded helical structure form those that remain manly disordered.

Based on these findings we next investigated to what extent typical IDP binding motifs promote pre-folded structures. We compiled a list of 20 IDP motifs from the literature and calculated the energetic contributions that stabilize helical structure (for their sequences, see Table S2). Some of these motifs also occur in the IDPs listed in the dataset (Table S1). The average helix propensity spans from −10 cal/mol to +280 cal/mol, with an average value of Δ*G*_CH_=+130 cal/mol. The average Δ*G*_CH_ value for IDP motifs is similar to the one we obtained for the contacting IDPs residues in our 65-IDP dataset, for which we have argued that prefolded structures are promoted (Fig. 4A). Additionally, we find that most of these motifs are also stabilized by *i-i+3* and *i-i+4* sidechain interactions, which on average add an additional −0.7 kcal/mol per motif. The resulting overall per-residue helix stabilization (sum of Δ*G*_CH_ and sidechain interactions) for IDP binding motifs is shown in Fig. 5B and listed in Table S2. The motifs conferring strongest stabilization of the helical conformation are of Vps27 and N-CoR ID2 with the X-X-a-ϕ-ϕ-X-X-ϕ-X-X-a-a and LXXI/HIXXXL/I motifs (Table S2). The presence of pre-folded structure in many of these motifs is also supported by the AGADIR calculation, when such motifs are embedded in the host IDP sequence (Fig. S11). Thus, many typical IDP binding motifs confer stability to helical elements in the unbound state, so promoting the formation of the pre-folded IDP structure.

## Discussion

We have systematical analyzed the extent of residual helical structure in intrinsically disordered proteins that fold to α-helix upon binding their targets. The average helix content is low, around 17%, and shows a rather uniform distribution (Fig. 2A). The location of helical regions in the unbound state correlates with their position in the target-bound state. In other words, the helical regions in unbound and bound state generally overlap and can be predicted with high accuracy (Fig. 2C). Our results support conclusions from a previous study, which established that IDP regions in the unbound state transiently sample the conformations that are present in their target bound state [13]. The study was based on the 24 IDP-target complexes that acquired different secondary structures, not only helical ones, and the overall correlation was around 60%. It therefore appears that the correlation between unbound and bound secondary structure is even stronger when only helical elements are considered (see Fig. 2C). More recently, a compilation of IDPs that form pre-structured motifs has been published, listing 47 mostly helical motif examples [11]. While the study established the presence of residual structure it did not analyze the overlap with the target-bound structure. The reported contents of the residual structure are somewhat higher, with IDPs exhibiting around 25% helicity on average. Collectively, our results show that pre-structured motifs are a rather general feature of α-helical binding motifs.

We next investigated the reasons governing the observed coupling between unbound and the target-bound helical structure. Using parameters describing the energetics of helix folding we have shown that the IDP residues that form contacts with the target have stronger propensity for helical conformation compared to the overall sequence or compared to non-contacting residues (which are part of the helical region in the target-bound conformation, but remain solvent exposed). The IDPs residues that form hotspots interactions with the target have the highest helix propensity (Fig. 4). We suggest that this can be, to a large extent, explained by the strong representation of leucine among hotspot residues, given that this residue has very helix propensity. It is well established that hydrophobic residues feature prominently in the protein-protein interactions. However, among them leucine has the highest helix propensity after alanine, while helix propensity of isoleucine is twofold lower and that of valine is threefold lower [35]. We have shown that a presence of leucine, but not other hydrophobic residues (isoleucine, valine) stabilizes pre-folded helical structure. Interestingly, the interfaces formed between globular proteins are depleted in hotspots mediated by leucine, and leucine is observed in less than 1% of hotspots, while its isomer isoleucine is much more common (observed in >9% of hotspots) [31]. It is tempting to speculate that the preference for leucine hotspots in the intrinsically disordered α-helical motifs is therefore related to the stabilization of pre-folded structure, that can be enabled by leucine but not for example isoleucine. In principle, the preformed motifs could be also stabilized by the non-contacting residues, for example by the presence of alanine at non-contacting positions, which is a common experimental strategy to artificially increase helix contents in IDPs [38]. This is however not the case, and the fraction of alanine is similar for contacting and non-contacting residues. In addition to the intrinsic helix propensity of amino acids we also analyzed the energetic contribution arising from the intrahelical sidechain interactions. In this case we observed that a significant number of such interactions are mediated by both contacting and non-contacting residues, although the nature of these sidechain interactions is different (Fig. S9). The energetic contributions to helix stabilization are approximately similar for both classes of residues, suggesting that in this case non-contacting residues also contribute to stabilization of pre-folded motifs via formation of electrostatic sidechain interactions. Collectively, the important result is that the pre-structured elements emerge due to presence of target-interacting residues with higher helix propensity, explaining why preformed-motifs are very common in helical IDPs. Thus, one may rephrase the previous observation that the unbound IDP structure predisposes the target-bound IDP conformation; rather the target-bound conformation is encoded by hotspot interactions that also define the unbound IDP structure.

The coupling between residual helicity and target-interactions is nicely illustrated by some common IDP α-helical binding motifs. For example, the binding motif LXXLL, observed in c-Myb and FOXO3a conserved region 2, stabilizes helical pre-folded structure due to high intrinsic helix propensity of leucine and the L-L sidechain interactions between *i-i+3* and *i-i+4* positions each conferring additional −0.23 and −0.5 kcal/mol, respectively (Fig. 5A, [34]). For a dataset of 20 such IDP motifs we have observed that many such motifs also stabilize the prefolded helical conformations (Fig. 5B). Interestingly, in some motifs we observed a combination of elements that both stabilize and destabilize helix conformation, possibly indicating that helix content, and related binding affinity, are modulated by evolution by including helix breakers. For example, the (L/V)(V/L)DGLL motif of the HBZ transcription factor that binds KIX domain contain many helix stabilizing features they also harbors a glycine residue at the solvent exposed side, which is a strong helix breaker [39]. It can be predicted that the presence of non-contacting glycine residue approximately halves the residual helicity of the (L/V)(V/L)DGL motif compared to one that would contain a helix-neutral non-contacting residue. Similarly, helix-breaking prolines are flanking the pre-structured motifs in the c-Myb and p57 proteins [15,17]. These cases might indicate that the residual helicity can be also finetuned by the presence of helix destabilizing elements.

In conclusion, our results indicate that pre-structured helical motifs are a general feature of disordered α-helical binding motifs. This is because IDP residues that mediate interactions with the target also tend to have high helix propensity. Thus, we suggest that it is the target-bound IDP structure that predisposes the pre-folded structure. The high occurrence of leucine, which has high helix propensity, among target-contacting and hotspot residues appears to be the main cause of this correlation between pre-folded and target-bound structure. Notably, the strong preference for leucine among IDP-target hotspots differs from what is observed in complexes between globular proteins, where leucine is underrepresented as a hotspot residue. The potential functional role of these pre-structured elements is still not clearly established. In several cases it has been shown that residual helicity is in fact functionally important [15,16] as it may tune the affinity of IDP-target interactions. However, even though simple thermodynamic arguments suggest that such pre-structuring would reduce the entropic penalty of folding and thereby increase the interaction affinity, more quantitative estimates of such stabilization are still lacking.

## Methods

### Residue definitions

Secondary structure and solvent accessible surface area (SASA) of target-bound IDPs was analyzed using STRIDE [40]. Contacting residues were defined as those that have buried side chains in the target-IDP complex. Accordingly contacting residue X had its SASA less than one third of the value for the tripeptide AXA minus 35 Å^2^ (AXA gives SASA for residue X in the tripeptide, 35 Å^2^ is area of main chain). In contrast, the non-contacting IDP residues were defined as those having SASA values higher than 30%, and could be either part of the helix or not linked to any kind of secondary structures (see text). Hotspot residues are a subset of contacting residues that establish energetically most favorable interactions. They were identified using a computational tool PPCheck [41]. Recently, we used several different tools to identify hotspot residues in IDP-target complexes such as alanine scanning methods (FoldX and Robbeta) and knowledge base tools (PPCheck and KFC). We observed that these tools give very similar hotspot predictions and largely agree with the experimental data, where available. The most consistent predictions were obtained by PPCheck, which is the reason why we used only this tool here.

### Circular dichroism spectroscopy and calculation of average helicity

In order to evaluate the alpha helical prediction of monomeric peptides made by AGADIR, experimental data (CD spectra) of 53 peptides, from IDPs that fold upon target binding, were obtained from literature. These spectra were digitalized (WebPlotDigitizer by Ankit Rohatgi) to extract the CD intensity at 222 nm. Additionally, 12 CD spectra were obtained in our laboratory (Figure S1). CD spectra were measured in 20mM phosphate buffer (150mM NaCl) pH 7 using JASCO J-1500 spectrometer. All peptides were diluted to the final concentration of 40μM as determined by UV spectrophotometry measuring absorbance at 280 nm. CD spectra were recorded in a cuvette with 1 mm optical path length at 25°C. CD spectrometer was calibrated prior measurements using ammonium-d-camphor-10-sulfonate (Sigma-Aldrich). The α-helical content of peptides in the dataset was determined from the mean residue molar ellipticity at 222 nm, using the equations from Errington et al. [20]. The dataset included the peptide sequence and the experimental parameters of the CD measurements (temperature, buffer pH and ionic strength). Possible interaction partners and the helicity in the bound state were determined using the Advanced Sequence search function of PDB.

### Estimation of conformation from NMR chemical shifts

The BMRB entries as reported in Table S1 were individually downloaded, and are available in the data/bmrb/ directory. BMRB entry 2426 was excluded at this stage as it contains only RDC values, resulting in 19 valid BMRB entries mapping to 20 peptide entries. The δ2D software [27] was executed on each BMRB entry using an in-house script, with the results saved per BMRB entry in files with .d2d suffix in the data/d2d/ directory. The cheSPI software [28] was executed on each BMRB entry using an in-house script, with the results saved in directories per BMRB entry in the data/cheSPI/ directory.

### Prediction from sequence

The peptide sequences used in CD measurements were analyzed with AGADIR [23] to determine its per-residues helical content by specifying temperature, buffer pH and ionic strength as used to acquire CD spectra. Using the b2bTools software [24], the backbone and sidechain dynamics, early folding propensity, and sheet, helix, coil and polyproline II propensities were predicted from sequence, all available from the *data/preds.json* file. Estimation of energetic contributions related to helix folding was performed by accounting for the intrinsic helicity contribution of each residues (Δ*G*_CH_) and the stabilization by sidechain interactions. The Δ*G*_CH_ contributions for each residue were summed together and divided by total number of residues. The Δ*G*_CH_ values form Pace and Sholtz were used and the Δ*G*_CH_=- 250 cal/mol for alanine was used [35,42]. The energetic contributions of sidechain interactions listed in the paper of Doig were used [34].

### Data integration

Using the custom *analysis.py* Python script (provided with datasets), the protein identifiers, information on the CD-determined experimental and AGADIR helicity for the full peptide, and the related BMRB entry (if available) were extracted using the Python pandas module from a the *data/final_helixpeptides.xlsx* Excel file, with the protein identifiers matched to the flat file (see next line). The per-residue AGADIR predicted helicity, final conformational state in the bound form (as determined by STRIDE), and information on the residues directly binding the target protein were extracted from the flat *data/final_ AGA_PP_STR.txt* and the *data/final_AGA_PP_STR_seqsForStrideData.txt* files containing this information with the in-script parseAgadir() function. The peptide sequence (CD determined in solution) was manually mapped to the bound peptide sequence. Entries without secondary structure information for the bound form (AFF4-3, AMBRA1, ATG16L1, ERM AAD) were excluded from the analysis, as well as minor sequence inconsistencies present. For these mismatched residues, no data was included in the further analysis.

## Acknowledgements

UZ acknowledges support through the “Young Researchers” program of the Slovenian Research Agency. This work was supported by the grants P1-0201 and J1-1706 from the Slovenian Research Agency to SH. WV acknowledges the Research Foundation Flanders (FWO) grant G.0328.16N.

## Data availability

The underlying data and analysis script is currently available for review from https://vubmy.sharepoint.com/:u:/g/personal/wim_vranken_vub_be/EY26ytACM3pBq3CPFXSPg3MBJ_R9aKD1v7EQw3FQtMWPwSQ?e=C5gJ48. Upon acceptance it will be made publicly available with a persistent DOI>

## Notes

### Competing Interest Statement

The authors have declared no competing interest.

