## Supporting information for "What stabilizes pre-folded structures in the intrinsically disordered α-helical binding motifs?"

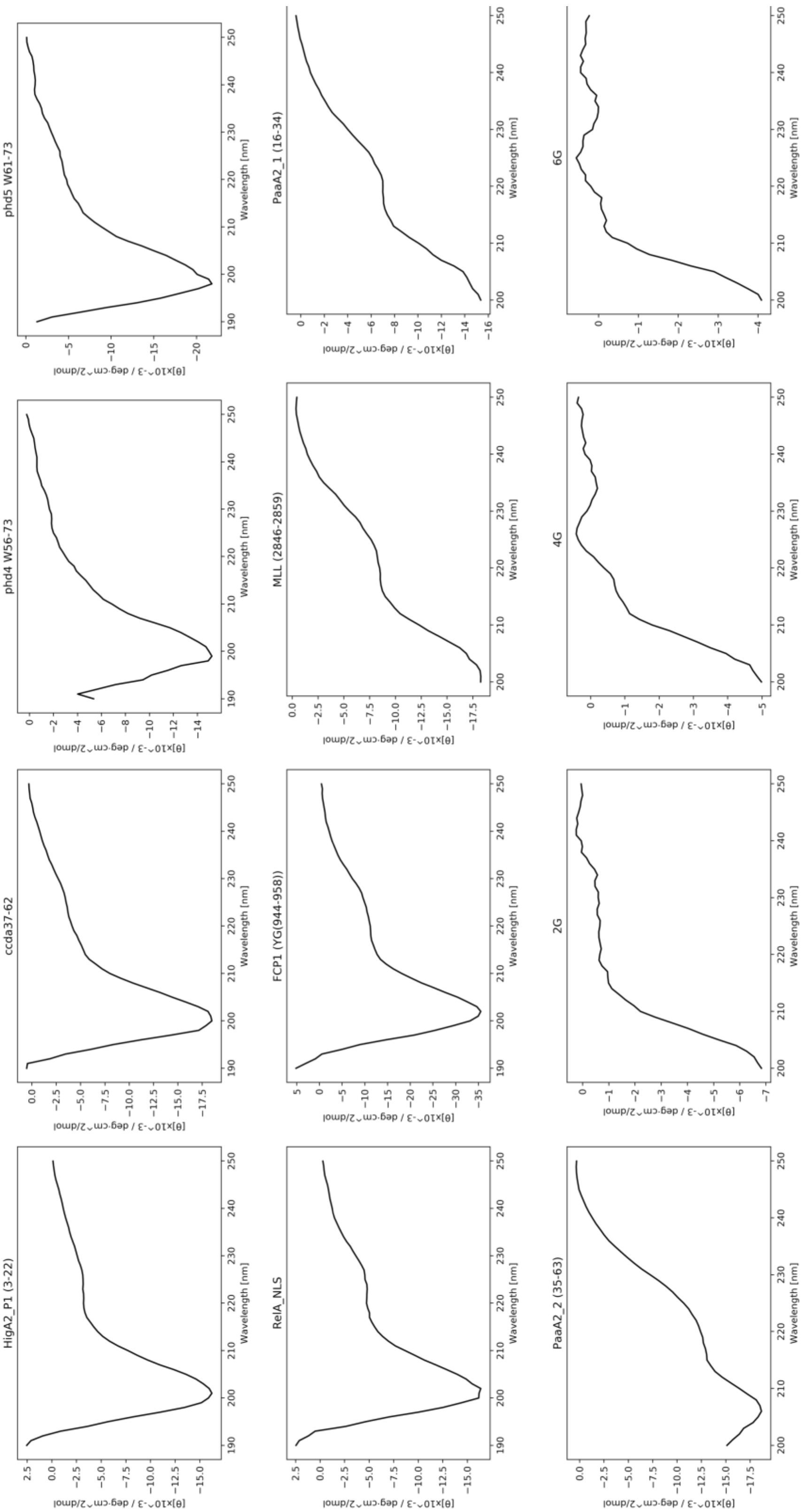

**Figure S1.** The CD spectra for a set of 12 peptides analyzed in our laboratory. CD spectra were measured as described in *Methods*. Peptide sequences are listed in Table S1.

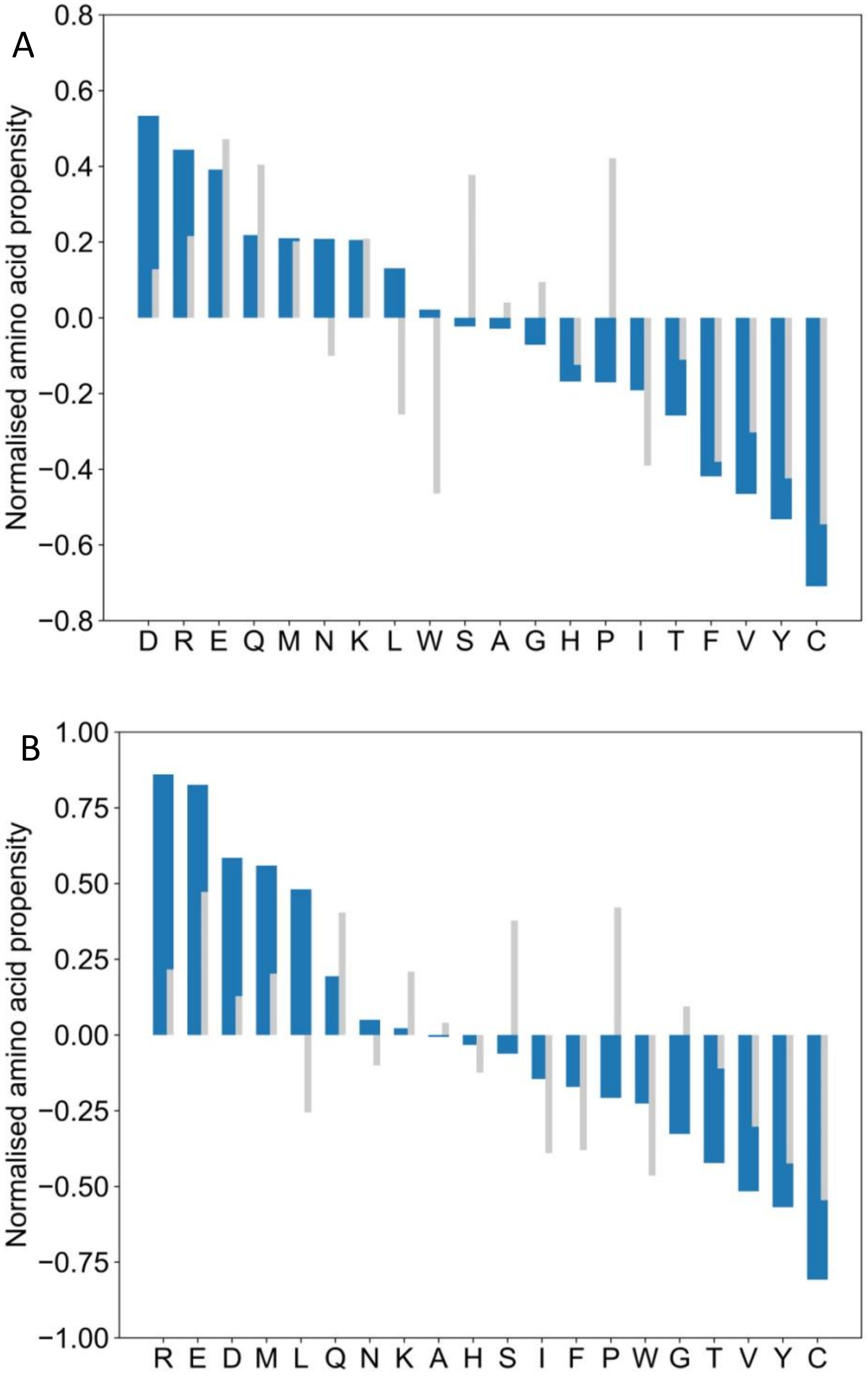

**Figure S2.** Comparison between the amino acid compositions of α-helical binding motifs from the reported dataset calculated using two different globular protein datasets. The amino acid frequencies of the globular proteins are either from Triphati et al. (panel A) or Mészáros et al. (panel B). Gray bars show the amino acid composition of IDPs from Disprot database.

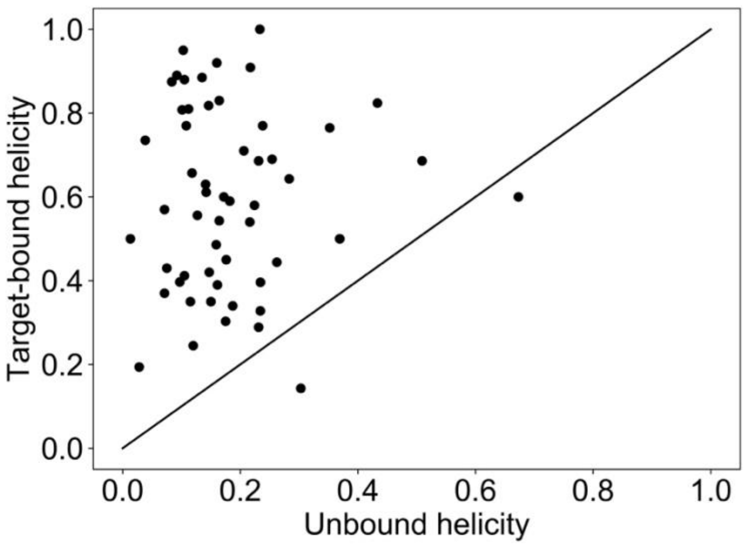

**Figure S3.** Helicity values for unbound and target-bound IDPs are not correlated. Data is shown in symbols and is taken form Table S1. Straight line with slope 1 and intercept 0 is shown for comparison.

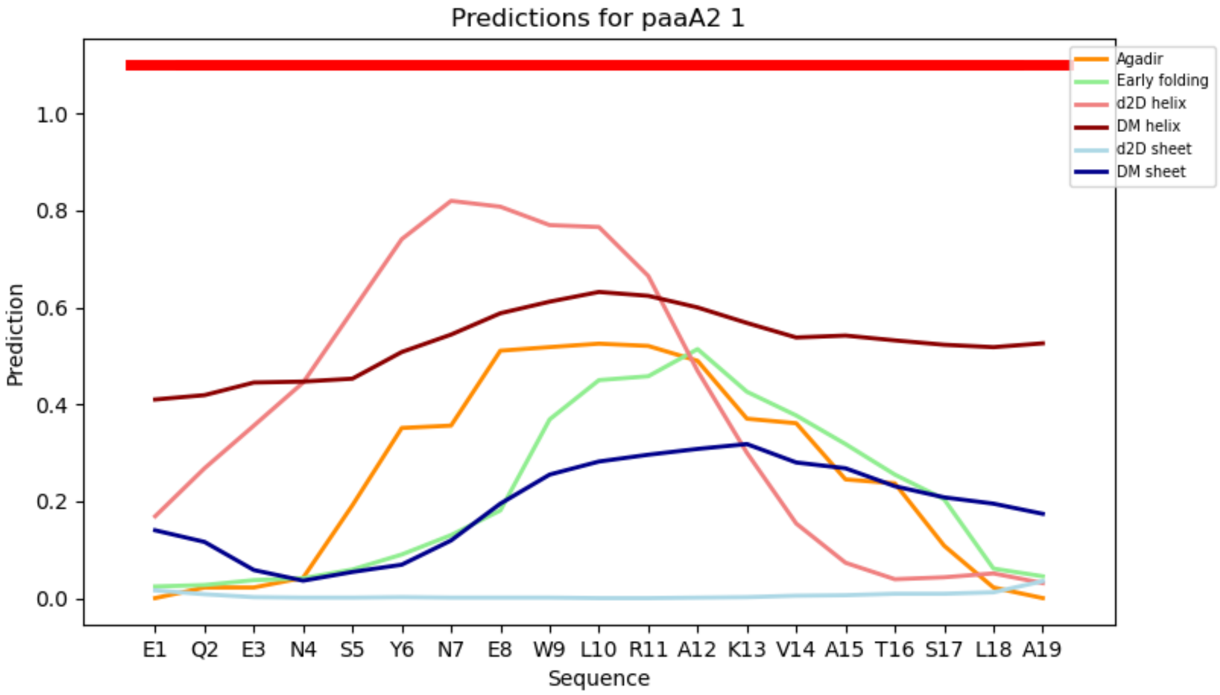

**Figure S4. Prediction of local IDP helicity using different sequence-based and chemical-shift based methods.** Lines in different colors show per-residue helicity as predicted by different methods. Thick red line above shows helical residues as observed in the target-bound IDP conformation. DM in legend text indicates DynaMine prediction.

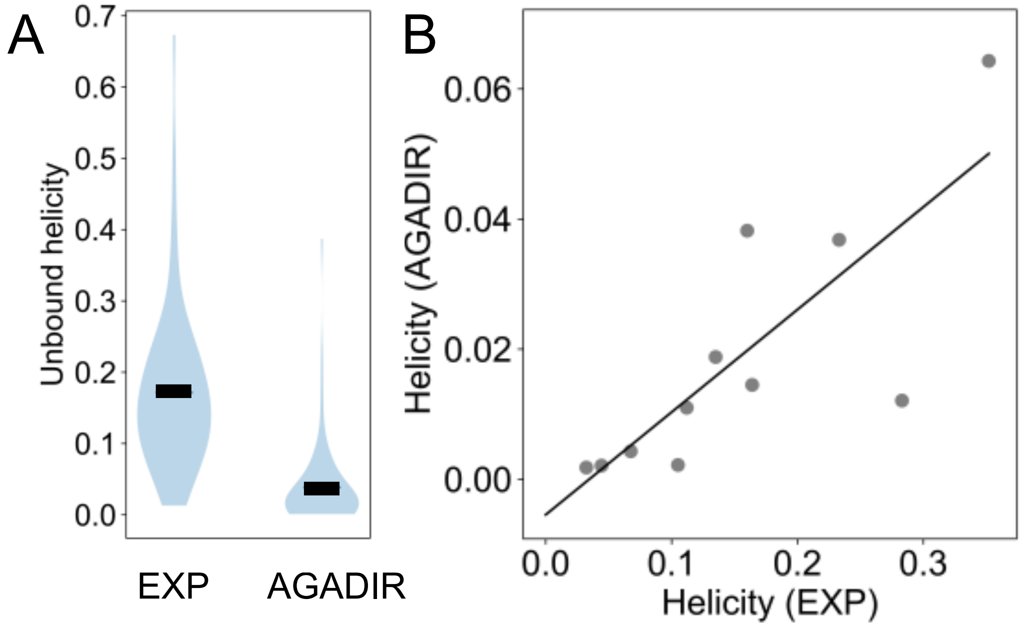

**Figure S5. A.** The average IDP helicity of α-helical binding motifs predicted by AGADIR is significantly lower compared to the experimental values from CD (t-test, $N_1$=$N_2$=65, $p$<$10^{-5}$). **B.** The experimental helicities for peptides measured in our laboratory correlate with those predicted by AGADIR, however the slope of the line is only 0.16 indicating that AGADIR underestimates absolute IDP helicity (R=0.79).

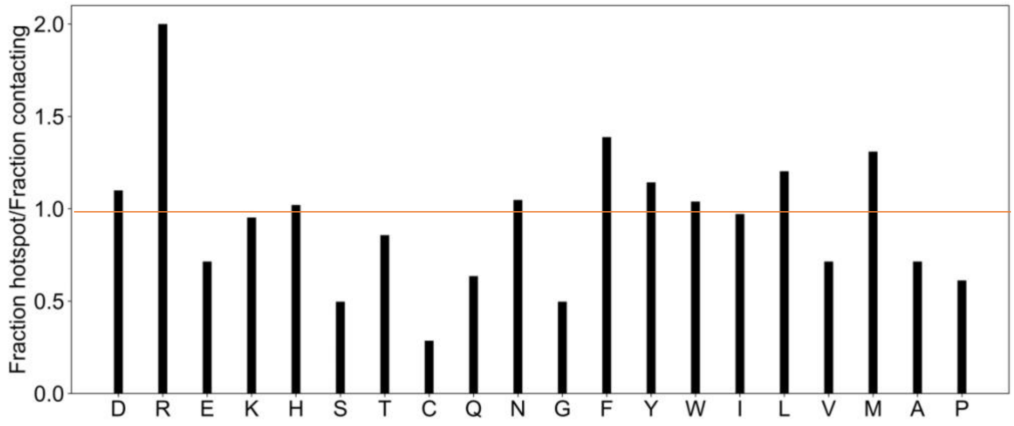

**Figure S6. Preference to form hotspots.** Bars show the ratio between hotspot frequency and contact frequency for each residue. In this way the hotspot fractions are normalized by the residue propensity to from any contact with the target. Values greater than 1 then indicate that the residue is more likely to form a hotspot when it establishes a contact with the target.

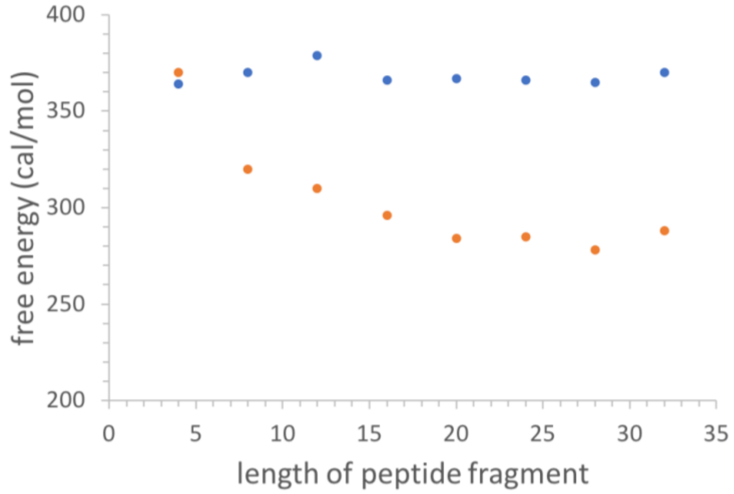

**Figure S7. Average energetic contributions for helix formation for peptides in Disprot.** We select 2000 random samples of peptides of different lengths (8-32) from the sequences listed in Disprot database and calculate their average (per-residue) intrinsic helix propensity $\Delta G_{CH}$ (blue circles) as well as the overall helix propensity (intrinsic + side-chain contributions) (shown in orange circles). Since interhelical side-chain interactions form between i-i+3 or i-i+4 residues, they depend on the length of peptide fragment, but this dependence levels off for peptides longer than 20 residues.

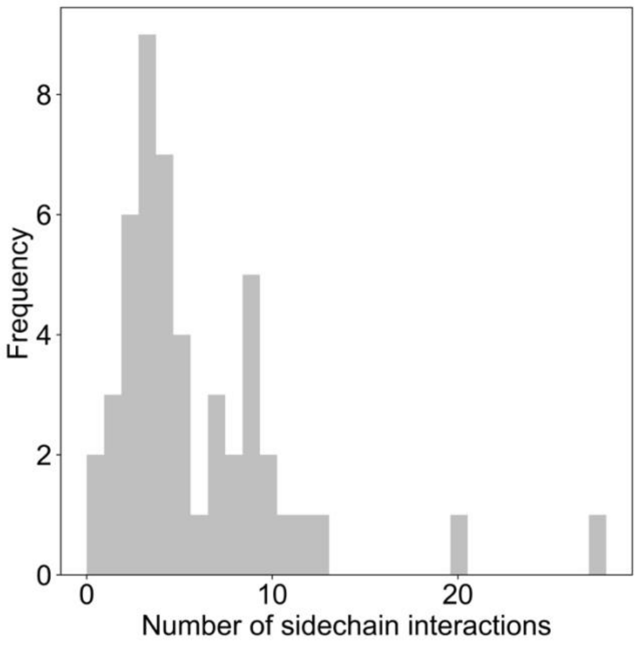

**Figure S8. Number of sidechain interactions contributing to helix stability.** The average number of identified sidechain interactions that affect helix energetics. Each IDP sequence was analyzed for different i-i+3 and i-i+4 sidechain interactions as listed in the paper by Doig (main text reference 34). The histogram shows the total number of such interactions obtained by analysis of all IDP residues (contacting and noncontracting).

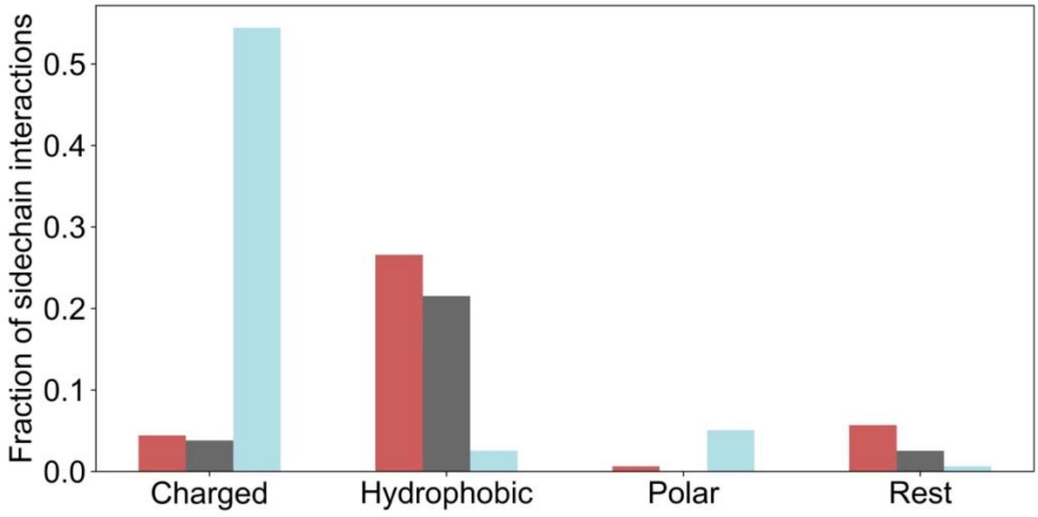

**Figure S9: Types of intrahelical sidechain interactions stabilizing helical structure.** Bars show the frequency of sidechain interactions between charged-charged (eg. E-K), hydrophobic-hydrophobic/aromatic (eg. L-L), polar-polar/charged (eg. Q-N), and other types of interactions (rest, these are mainly charged-aromatic and charged-hydrophobic interactions (eg. F-K). These sidechain interactions form between residues at position i and at position i+3 or i+4. Colors denote that both residues classified as target-contacting (red), hotspots (gray) or non-contacting (blue).

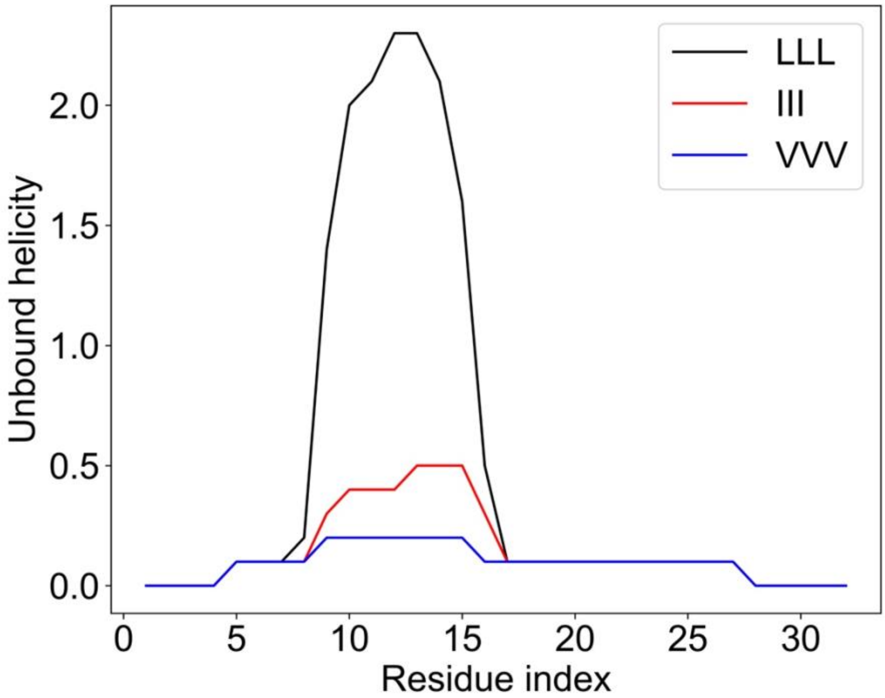

**Figure S10: Leucine hotspots, but not isoleucine or valine motifs stabilize pre-folded helices.** AGADIR calculated helicity (in %) of unbound model IDP shows the effects of leucine, isoleucine and valine. It is evident that preferential use of leucine in IDP-target hotspots, but not isoleucine or valine, will lead to considerable stabilization of pre-folded helical structure.

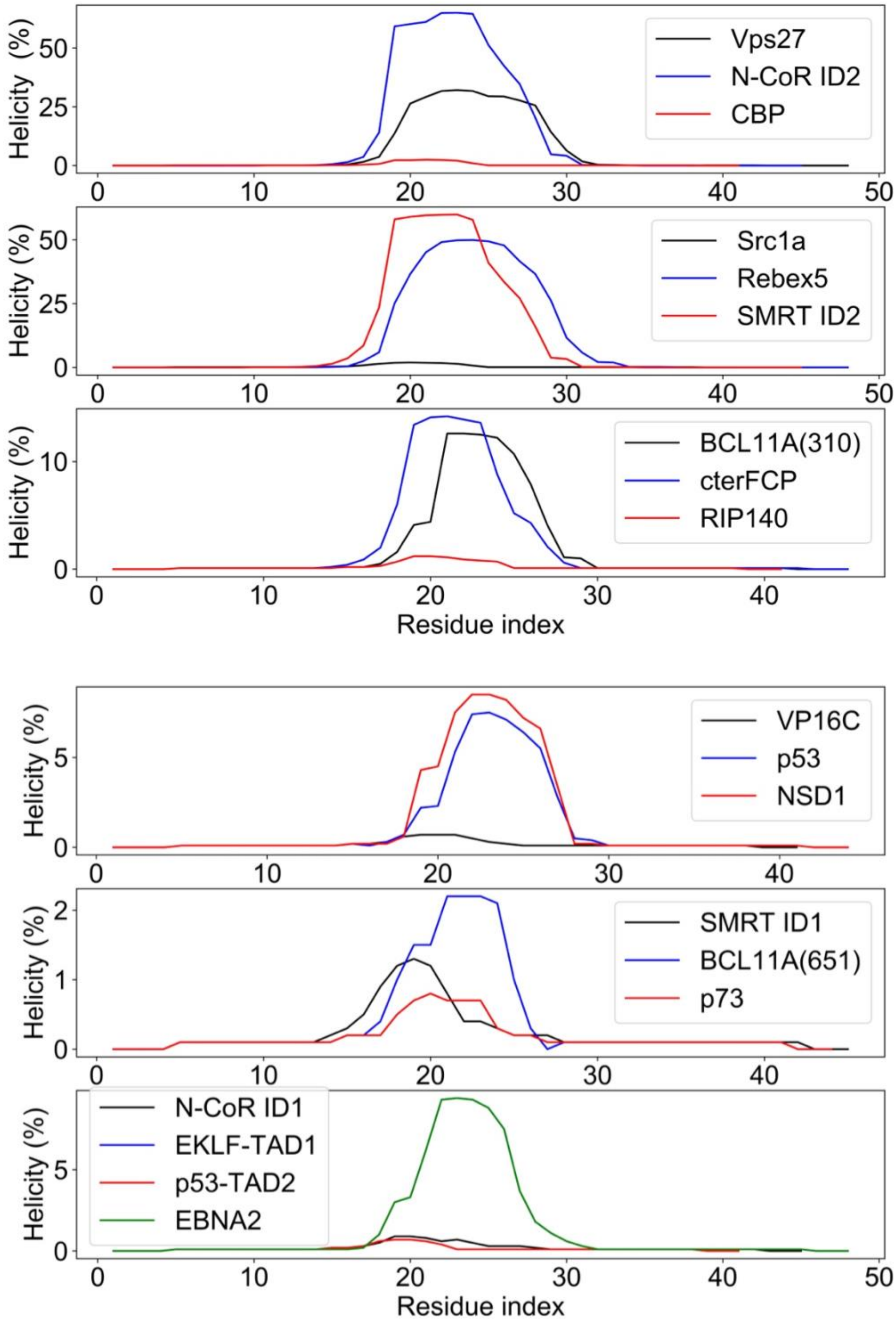

**Figure S11. AGADIR calculated helicity for intrinsically disordered α-helical binding motifs show pre-folded structures.** Each characterized IDP binding motif (listed in Table S3) was inserted into the host sequence: $(SETSTTSSS)_2$(motif)$(SETSTTSSS)_2$. The host sequence itself has the same average helix propensity as a random sample obtained from Disprot database.

**Table S1. A database of experimental helicities of the α-helical recognition elements.** Symbol I denotes ionic strength, while T is temperature in ° C.

| IDP | UNIPROT IDP | SEGMENT | length | conditions (T/pH/I) | Helicity (exp,CD) | BMRB entry | bindign partenr | UNIPROT target | PDB | $\alpha_H$ | ΔHelicity | reference |
|---|---|---|---|---|---|---|---|---|---|---|---|---|
| calcineurin | Q08209 | 396-414 | 19 | 5/7/0,1 | 0.103 | 26990 | calmodulin | P0DP23 | 4Q5U | 0.950 | 0.847 | Peptides: Frontiers of Peptide Science |
| ACTR 2 | Q9Y6Q9 | 1046-1058 | 13 | 5/7/0,1 | 0.238 | 15397 | CBP | P45481 | 1KBH | 0.770 | 0.532 | https://www.ncbi.nlm.nih.gov/pubmed/20556825 |
| ACTR 3 | Q9Y6Q9 | 1073-1085 | 13 | 5/7/0,1 | 0.108 | 15397 | CBP | P45481 | 1KBH | 0.770 | 0.662 | https://www.ncbi.nlm.nih.gov/pubmed/20556825 |
| Hif-1α CAD | Q16665 | 776-826 | 51 | 25/6,9/0,1 | 0.075 | | CBP (TAZ1) | P45481 | 1L8C | 0.430 | 0.355 | https://www.nature.com/articles/s41598-018-26213-x |
| CITED2 | Q99967 | 220-269 | 50 | 25/7,5/0,1 | 0.187 | | CBP (TAZ1) | P45481 | 1R8U | 0.340 | 0.153 | https://www.ncbi.nlm.nih.gov/pubmed/14594809 |
| p21 | P38936 | 9-84 | 76 | 5/7/0,2 | 0.231 | 17427 | CDK4 | P11802 | 6P8H | 0.289 | 0.058 | https://www.pnas.org/content/pnas/93/21/11504.full.pdf |
| STAT2 | P52630 | 786-838 | 53 | 25/6,9/0,1 | 0.115 | | CBP (TAZ1) | P45481 | 2KA4 | 0.350 | 0.235 | https://pubs.acs.org/doi/pdf/10.1021/acs.biochem.7b00428?rand=m |
| PUMA | Q99ML1 | 127-161 | 35 | 5/7/0,1 | 0.231 | | Mcl-1 | P97287 | 2ROC | 0.686 | 0.455 | http://www.jbc.org/content/suppl/2018/05/01/RA118.002791.DC1/13 |
| BID | P70444 | 76-110 | 35 | 5/7/0,1 | 0.172 | | Bcl2-A1 | Q07440 | 2VOI | 0.600 | 0.428 | http://www.jbc.org/content/suppl/2018/05/01/RA118.002791.DC1/13 |
| RelB2 | P0C079 | 47-79 | 33 | 25/7/0,1 | 0.224 | 16067 | RelE mRNA interferaza | P0C077 | 4FXE | 0.580 | 0.356 | https://www.sciencedirect.com/science/article/pii/S00222836080047 |
| TPX2 | Q9ULW0 | 30-43 | 14 | 25/7/0,1 | 0.071 | | Aurora-A | O14965 | 1OL5 | 0.570 | 0.499 | https://www.ncbi.nlm.nih.gov/pubmed/27775325 |
| WASP | P42768 | 221-257 | 37 | 25/6.5/0.1 | 0.082 | | CDC42 | P60953 | 1CEE | | | https://www.jbc.org/content/273/29/18067.full#sec-1 |
| VP35 | Q05127 | 28-48 | 21 | 5/7/0,1 | 0.303 | | Nucleoprotein | P18272 | 4YPI | 0.143 | - 0.160 | https://www.cell.com/cell-reports/fulltext/S2211-1247(15)00303-4?_r |
| TRAP220 | Q15648 | 637-654 | 18 | 4/7,4/0,1 | 0.262 | | PPARgamma LBD | P37231 | 6ONJ | 0.444 | 0.182 | https://www.ncbi.nlm.nih.gov/pubmed/27179590 |
| BECN1 | Q14457 | 105-130 | 26 | 4/7,4/0,1 | 0.101 | 25384 | Bcl-2-L1 | Q07817 | 2P1L | 0.808 | 0.707 | https://www.ncbi.nlm.nih.gov/pubmed/27179590 |
| p53 | P04637 | 367-388 | 22 | 4/7,4/0,1 | 0.013 | | S100B | P04631 | 1DT7 | 0.500 | 0.487 | https://www.ncbi.nlm.nih.gov/pubmed/27179590 |
| AMBRA1 | Q9C0C7 | 527-541 | 15 | 4/7,4/0,1 | 0.208 | | | | | | | https://www.ncbi.nlm.nih.gov/pubmed/27179590 |
| ATG16L1 | Q676U5 | 59-84 | 26 | 4/7,4/0,1 | 0.093 | | | | | | | https://www.ncbi.nlm.nih.gov/pubmed/27179590 |
| p27ID | P46527 | 22-97 | 76 | 5/7,5/0,1 | 0.175 | 6112 | CDK2, CCNA2 | P24941 | 1JSU | 0.303 | 0.128 | https://www.ncbi.nlm.nih.gov/pubmed/11790096 |
| p65 | Q04206 | 522-551 | 30 | 20/7/0,1 | 0.071 | | CBP (KIX) | P45481 | 5U4K | 0.370 | 0.299 | http://www.jbc.org/content/269/41/25613.long |
| NTAIL | Q77M43 | 486-505 | 20 | 20/7/0,1 | 0.092 | 6566 | virus P protein | Q77M42 | 1T6O | 0.890 | 0.798 | https://www.ncbi.nlm.nih.gov/pubmed/29197511 |
| E2A | P15884 | 9-27 | 19 | 10/7,4/0,1 | 0.141 | 50196 | CBP (KIX) | Q92793 | 2KWF | 0.630 | 0.489 | https://www.pnas.org/content/111/33/12055 |
| SRC1 (NCOA1) | Q15788 | 920-970 | 51 | 25/7,4/0,1 | 0.161 | | CBP (NCBD) | P45481 | 2C52 | 0.390 | 0.229 | https://www.ncbi.nlm.nih.gov/pubmed/26153298 |
| TRTK12 | P13127 | 265-276 | 12 | 25/7,5/0,1 | 0.147 | | protein 16A | P14315, Q6EDY6 | 3AA0 | 0.420 | 0.273 | https://pubs.acs.org/doi/10.1021/bi300865g |
| NDR | Q15208 | 62-87 | 26 | 25/7,5/0,1 | 0.216 | | S100B | P02638 | 1PSB | 0.540 | 0.324 | https://pubs.acs.org/doi/10.1021/bi300865g |
| HDM2 | Q00987 | 25-44 | 20 | 25/7,5/0,1 | 0.133 | 15945 | | | | | | https://pubs.acs.org/doi/10.1021/bi300865g |
| HDM4 | O15151 | 25-44 | 20 | 25/7,5/0,1 | 0.178 | 25546 | | - | | | | https://pubs.acs.org/doi/10.1021/bi300865g |
| $IA_3$ | P01094 | 1-68 | 68 | 25/7/0,1 | 0.097 | 6078 | Proteinaza A | P07267 | 1DPJ | 0.397 | 0.300 | https://www.ncbi.nlm.nih.gov/pubmed/15065849 |
| pKID | P15337 | 102-132 | 31 | 10/7.4/0.1 | 0.206 | 6784 | CBP (KIX) | P45481 | 1KDX | 0.710 | 0.504 | https://www.pnas.org/content/111/33/12055 |

| | | | | | | | | | | | | |
|---|---|---|---|---|---|---|---|---|---|---|---|---|
| MLL | Q03164 | 2838-2869 | 32 | 10/7.4/0.1 | 0.150 | | CBP + Myb | P06876, P45481 | 2AGH | 0.350 | 0.200 | https://www.pnas.org/content/111/33/12055 |
| FOXO3a:CR2C | O43524 | 462-483 | 22 | 20/6/0.1 | 0.182 | | CBP | P45481 | 2LQH | 0.590 | 0.408 | https://www.pnas.org/content/pnas/109/16/6078.full.pdf |
| FOXO3a:CR3 | O43524 | 620-635 | 16 | 20/6/0.1 | 0.254 | | CBP | P45481 | 2LQH | 0.690 | 0.436 | https://www.pnas.org/content/pnas/109/16/6078.full.pdf |
| FITC-cMyb | P06876 | 291-315 | 25 | 25/7.4/0.1 | 0.217 | | CBP + HBZ | P45481, Q2Q067 | 6DNQ | 0.909 | 0.692 | https://www.ncbi.nlm.nih.gov/pubmed/23875714 |
| S peptide | P61823 | 27-46 | 20 | 1.0/4.03/0.2 | 0.176 | 2426 | S protein | P61823 | 1CJR | 0.450 | 0.274 | https://www.sciencedirect.com/science/article/pii/002228368290168 |
| BAK | O08734 | 62-96 | 35 | 25/7/0.1 | 0.159 | | BCL2L1 | Q07817 | 5FMJ | 0.486 | 0.326 | https://www.sciencedirect.com/science/article/abs/pii/S00222836183 |
| BAX | Q07813 | 49-83 | 35 | 25/7/0.1 | 0.118 | 4632 | BCL2 | P10415 | 2XA0 | 0.657 | 0.539 | https://www.sciencedirect.com/science/article/abs/pii/S00222836183 |
| BIM | O54918 | 136-170 | 35 | 25/7/0.1 | 0.146 | | BCL2L1 | Q64373 | 1PQ1 | 0.818 | 0.672 | https://www.sciencedirect.com/science/article/abs/pii/S00222836183 |
| BMF | Q91ZE9 | 124-158 | 35 | 25/7/0.1 | 0.673 | | BCL2A1 | Q07440 | 2VOG | 0.600 | | https://www.sciencedirect.com/science/article/abs/pii/S00222836183 |
| NOXA_A | Q9JM54 | 13-47 | 35 | 25/7/0.1 | 0.164 | | MCL-1 | P97287 | 2ROD | 0.543 | 0.379 | https://www.sciencedirect.com/science/article/abs/pii/S00222836183 |
| NOXA_B | Q9JM54 | 64-98 | 35 | 25/7/0.1 | 0.509 | | MCL-1 | P97287 | 2JM6 | 0.686 | 0.177 | https://www.sciencedirect.com/science/article/abs/pii/S00222836183 |
| DREB2A | O82132 | 255-272 | 18 | 25/7/0.1 | 0.127 | 34152 | RCD1-RST | - | 5OAP | 0.556 | 0.429 | https://www.sciencedirect.com/science/article/pii/S09692126183009 |
| ERM AAD | P41161 | 43-72 | 34 | 20/10.6/0.1 | 0.102 | | | | 4UNO | | | https://academic.oup.com/nar/article/25/22/4455/2359278 |
| CeRED (Smu-2) | Q9N4U5 | 163-223 | 61 | 20/8/0.1 | 0.234 | | SMU-1 | G5EEG7 | 5EN7 | 0.328 | 0.094 | https://www.cell.com/structure/fulltext/S0969-2126(16)00097-6?_ret |
| AFF4-1 | Q9UHB7 | 2-73 | 72 | 25/7.5/0.1 | 0.028 | | CDK, Tat, Cyclin-T1 | P50750 | 4OR5 | 0.194 | 0.167 | https://www.pnas.org/content/110/2/E123 |
| AFF4-3 | Q9UHB7 | 710-729 | 20 | 25/7.5/0.1 | 0.024 | | | | | | | https://www.pnas.org/content/110/2/E123 |
| HigA2 | Q9KMA5 | 3-23 | 21 | 25/7.4/0.17 | 0.112 | | HigB-2 | Q9KMA6 | 5JAA | 0.810 | 0.698 | this study (Figure S1) |
| Ccda | P62552 | 37-62 | 26 | 25/7.4/0.17 | 0.135 | 6743 | ccdB | P62554 | 3HPW | 0.885 | 0.750 | this study (Figure S1) |
| Phd4 | Q06253 | 56-73 | 18 | 25/7.4/0.17 | 0.105 | | Doc | Q06259 | 3K33 | 0.880 | 0.775 | this study (Figure S1) |
| Phd5 | Q06253 | 61-73 | 13 | 25/7.4/0.17 | 0.164 | | Doc | Q06259 | 3K33 | 0.830 | 0.666 | this study (Figure S1) |
| RelA_NLS | Q04207 | 293-316 | 24 | 25/7.4/0.17 | 0.16 | 25792 | IkappaBbeta | Q60778 | 1K3Z | 0.920 | 0.760 | this study (Figure S1) |
| FCP1 | Q9Y5B0 | YG944-958 | 17 | 25/7.4/0.17 | 0.352 | 16296 | RAP74 | P35269 | 1J2X | 0.765 | 0.413 | this study (Figure S1) |
| MLL | Q0316 | 2846-2859 | 14 | 25/7.4/0.17 | 0.283 | | CBP | Q92793 | 2LXS | 0.643 | 0.360 | this study (Figure S1) |
| HigA22G* | | | 22 | 25/7.4/0.17 | 0.068 | | | | 5JAA | | | this study (Figure S1) |
| HigA24G* | | | 22 | 25/7.4/0.17 | 0.044 | | | | 5JAA | | | this study (Figure S1) |
| HigA26G* | | | 22 | 25/7.4/0.17 | 0.032 | | | | 5JAA | | | this study (Figure S1) |
| PaaA2 1 | Q8XAD5 | 16-34 | 19 | 25/7.4/0.17 | 0.233 | 18841 | ParE2 | A0A0H3JHG3 | 5CW7 | 1.000 | 0.767 | this study (Figure S1) |
| PaaA2 2 | Q8XAD5 | 35-63 | 30 | 25/7.4/0.17 | 0.369 | 18841 | ParE2 | A0A0H3JHG3 | 5CW7 | 0.500 | 0.131 | this study (Figure S1) |
| p53 | P04637 | 13-61 | 49 | - | 0.12 | 17760 | CBP | P45481 | 2L14 | 0.245 | 0.125 | https://pubs.acs.org/doi/10.1021/bi1012996#notes-1 |
| HIND-I | Q12420 | 7-24 | 18 | 25/7.6/0.1 | 0.142 | | HUB1 | Q6Q546 | 3PLU | 0.611 | 0.469 | https://www.nature.com/articles/nature10143#Sec22 |
| NCOA1 CID | Q15788 | 924-964 | 41 | 4/7.4/0.2 | 0.18 | | CBP | | 2C52 | | | https://prod--journal.elifesciences.org/articles/16059v2/figures |
| 4E-BP1 | Q13541 | 51-67 | 17 | 25/7.0/0 | 0.105 | 27003 | eIF4E2 | O60573 | 2JGB | 0.412 | 0.307 | https://pubmed.ncbi.nlm.nih.gov/10394359/ |
| SID peptide (MAD1) | Q05195 | 7-20 | 17 | 25/7/0.1 | 0.433 | | Sin3A | Q60520 | 1G1E | 0.824 | 0.391 | https://www.jbc.org/article/S0021-9258(17)46573-2/fulltext |
| HBZ_1 | Q2Q067 | 3-36 | 34 | 28/6.8/0.2 | 0.038 | 27517 | Myb + CBP | P06876 | 6DNQ | 0.735 | 0.697 | https://pubmed.ncbi.nlm.nih.gov/30232260/ |
| HBZ_2 | Q2Q067 | 33-56 | 24 | 28/6.8/0.2 | 0.084 | 27517 | Myb + CBP | P45481 | 6DNQ | 0.875 | 0.791 | https://pubmed.ncbi.nlm.nih.gov/30232260/ |
| c-myb | P06876 | 275-327 | 53 | 25/6.8/0.2 | 0.234 | | CBP | P45481 | 1SB0, 6DNQ | 0.396 | 0.162 | https://pubmed.ncbi.nlm.nih.gov/29890118/ |

*sequnces of HigA2 2G, 4G and 6G are: NRDLFGELSSALGEAKQHSEGW, NRDLFGELSGALGEAKGHSEGW, NRDLFGELGGALGEAKGHGEGW

**Table S2**: Individual performances of predictors in identifying residues that fold into a helical conformation when binding. The closer the value to 1.0, the better the prediction. Values under 0.5 mean the value is negatively correlated, and therefore is equivalent to an AUC of 1.0 minus that value.

| **Predictor – sequence based** | ROC AUC |
|---|---|
| AGADIR | 0.81 |
| DynaMine backbone rigidity | 0.68 |
| DynaMine coil propensity | 0.28 (reverse AUC 0.72) |
| DynaMine helix propensity | 0.78 |
| DynaMine sheet propensity | 0.48 |
| EFoldMine early folding propensity | 0.75 |
| **Estimation – chemical shift based** | |
| δ2D coil | 0.55 |
| δ2D helix | 0.63 |
| δ2D sheet | 0.21 (reverse AUC 0.79) |
| cheSPI coil | 0.35 (reverse AUC 0.65) |
| cheSPI helix | 0.70 |
| cheSPI sheet | 0.70 |

**Table S3.** List of IDP binding motifs. X - any residue, a - an acidic residue (Asp or Glu), ϕ - bulky hydrophobic residue. Last three columns list the energetic contributions related to helix stability arising from the intrinsic helix propensity (intrinsic, in cal/mol per residue), from the sidechain interactions (sidechain, in cal/mol per motif) and sum of both (total, in cal/mol per residue).

| **IDP** | **motif** | **sequence** | **total** | **intrinsic** | **sidechain** |
|---|---|---|---|---|---|
| Vps27 (259-270) | X-X-a-ϕ-ϕ-X-X-ϕ-X-X-a-a | SLEIAKRILEEE | -109 | 69 | -2140 |
| N-CoR ID2 | LXXI/HIXXXL/I | LEDIIRKAL | -104 | 63 | -1500 |
| CBP | LXXLL | LSELL | -96 | 58 | -770 |
| Src1a | LXXLL | LVQLL | -76 | 78 | -770 |
| Rabex5 (54-65) | X-X-a-ϕ-ϕ-X-X-ϕ-X-X-a-a | DWELAERLQREE | -74 | 86 | -1920 |
| SMRT ID2 | LXXI/HIXXXL/I | LEAIIRKAL | -65 | -14 | -460 |
| BCL11A (310) | F/YSXXLXXL/Y | FSRRLREL | -42 | 63 | -840 |
| cterFCP | L/IEXELXDhh | LEAELNDLM | 1 | 86 | -770 |
| RIP140 | LXXLL | LEGLL | 4 | 158 | -770 |
| VP16C | ϕXXϕϕ | FEQMF | 14 | 174 | -800 |
| VP16 | ϕXXϕϕ | FEQMF | 14 | 174 | -800 |
| p53-TAD1 | FXXϕWXXL | FSDLWKLL | 29 | 140 | -890 |
| NSD1 | F/YSXXLXXL/Y | FSTLLMML | 35 | 103 | -540 |
| SMRT ID1 | LXXI/HIXXXL/I | LAQHISEVI | 55 | 140 | -760 |

| | | | | | |
|---|---|---|---|---|---|
| BCL11A (651) | F/YSXXLXXL/Y | YSQWLAGY | 98 | 208 | -880 |
| p73-TAD | FXXϕWXXL | FEHLWSSL | 110 | 178 | -540 |
| N-CoR ID1 | LXXI/HIXXXL/I | LADHICQII | 170 | 170 | 0 |
| EKLFTAD1 (24-35) | X-X-a-ϕ-ϕ-X-X-ϕ-X-X-a-a | QDDFLKWWRSEE | 175 | 191 | -190 |
| p53-TAD2 | ϕXXϕϕ | IEQWF | 198 | 198 | 0 |
| p53 | ϕXXϕϕ | IEQWF | 198 | 198 | 0 |
| EBNA2 | ϕXXϕϕ | WDYIF | 284 | 284 | 0 |
